# Combining auxin-induced degradation and RNAi screening identifies novel genes involved in lipid bilayer stress sensing in *Caenorhabditis elegans*

**DOI:** 10.1101/2020.05.20.107151

**Authors:** Richard Venz, Anastasiia Korosteleva, Collin Y. Ewald

## Abstract

Alteration of the lipid composition of biological membranes interferes with their function and can cause tissue damage by triggering apoptosis. Upon lipid bilayer stress, the endoplasmic reticulum mounts a stress response that is similar to the unfolded protein response. However, only a few genes are known to regulate lipid bilayer stress. Here, we performed a suppressor screen that combined the auxin-inducible degradation (AID) system with conventional RNAi in *C. elegans* to identify members of the lipid bilayer stress response. AID-mediated knockdown of the mediator MDT-15, a protein required for the upregulation of fatty acid desaturases, caused activation of a lipid bilayer stress sensitive reporters. We screened through almost all *C. elegans* kinases and transcription factors using RNAi by feeding. We report the identification of 8 genes that have not been implicated previously with lipid bilayer stress before in *C. elegans*. These suppressor genes include *skn-1*/NRF1,2,3 and *let-607*/CREB3. Our candidate suppressor genes suggest a network of transcription factors and the integration of multiple tissues for a centralized lipotoxicity response in the intestine. Additionally, we propose and demonstrate the proof-of-concept for combining AID and RNAi as a new screening strategy.

## Introduction

Biological membranes play an important role in protein folding, signaling, secretion, and turnover of proteins. Changes in the lipid composition of a membrane alters its properties, thus, interferes with its function and leads to lipid bilayer stress (LBS) (Covino et al., 2018). Maintaining the membranes’ composition is therefore crucial for a cell. High dietary intake of saturated fatty acids leads to a metabolic syndrome referred to as lipotoxicity (Ertunc and Hotamisligil, 2016). On a cellular level, elevated levels of saturated fatty acids alter membrane composition. Sensitive for these changes is the endoplasmic reticulum, which is a major site for protein and lipid synthesis, and the main intracellular calcium storage (Schwarz and Blower, 2016). Lipid disequilibrium interferes with secretory capacity, and renders cells specialized in secretion, such as insulin-producing beta cells, susceptible to cell death (Preston et al., 2009, Acosta-Montaño and García-González 2018). Although LBS has been suggested to play a major part in disease progression, the spectrum of the underlying molecular players sensing LBS remains to be identified.

The unfolded protein (UPR) sensors IRE1, PERK1, and ATF6 have been found to be sensitive for changes in membrane fluidity (Volmer et al., 2013; Koh et al., 2018). On the molecular level, IRE1, PERK1, and ATF6 act in parallel in response to unfolded proteins (Fig. 1a). Activated IRE1 splices *XBP1* mRNA to stabilize the transcript and to allow translation of the spliced XBP1 transcription factor (Fig. 1a). PERK1 phosphorylates the initiation factor eIF2alpha, which reduces translation rate and allows preferential translation of genes containing upstream open reading frames (uORFs), such as the transcription factor ATF4 (Harding et al., 2000; Fig. 1a). During ER stress, ATF6 translocates from the ER to the Golgi, where it is cleaved by proteases called S1P and S2P. Cleaved ATF6 migrates to the nucleus and acts as a transcription factor (Fig. 1a). The transcription factors co-regulate many targets, but how the downstream targets of the three arms of the UPR restore membrane homeostasis in detail remains unknown.

**Figure 1.**
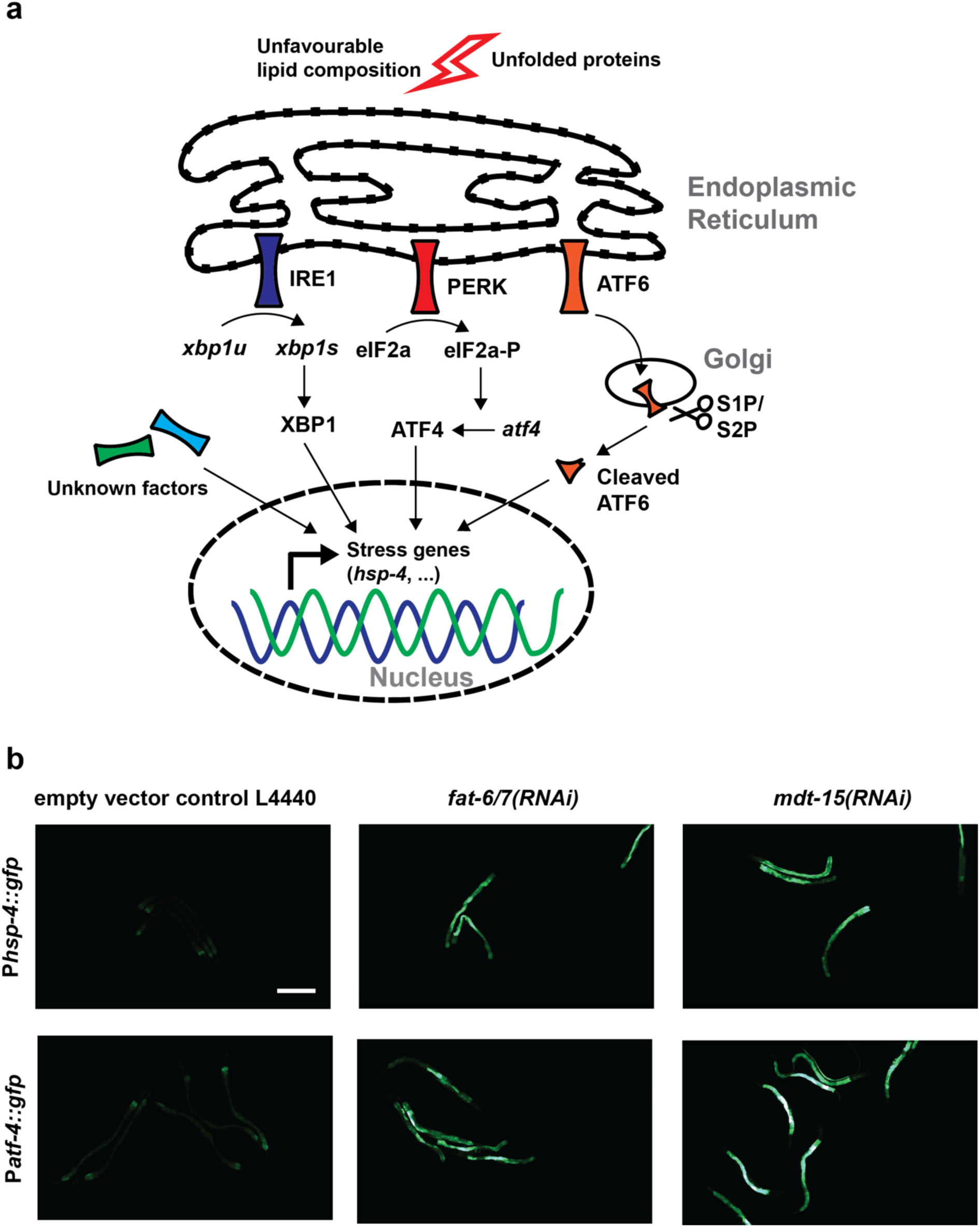
Integrated stress response of *C. elegans*. a) Model of the unfolded protein and lipid bilayer stress response. b) P*hsp-4::gfp* and P*atf-4::gfp* are activated by *fat-6/7(RNAi)* and *mdt-15(RNAi)*. Bar = 200 μm.

In *C. elegans*, loss of fatty acid desaturases *fat-6* and *fat-7* or the mediator *mdt-15*, which regulates *fat-6/7* expression, leads to higher ratio of saturated fatty acids in the membranes. This activates the ER stress reporter *hsp-4::gfp* via the IRE-1/XBP-1 axis (Hou et al., 2014). Activation of the ER stress sensor can also be achieved by depleting the cell’s phosphatidylcholine levels. Curiously, the signature of lipid bilayer stress response is different from the canonical UPR in *C. elegans* (Hou et al., 2014; Koh et al., 2018). This argues for an additional layer of regulation that fine-tunes the output during activation of the three UPR arms (Fig. 1a).

Genetic mutant screening for members of the UPR have been successful (Calfon et al., 2002; Singh and Aballay, 2017), but might have missed lethal genes. RNAi-based forward screens can bypass genes that cause embryonal lethality or developmental defects. However, feeding more than one RNAi simultaneously was previously reported to produce poor results (Min et al., 2010). This suggests a bottleneck for screening strategies where one would like to screen for suppressors of a phenotype caused by a knock down using an RNAi-mediated screen.

The auxin inducible degradation (AID) has been introduced into *C. elegans* recently (Zhang et al., 2015). A protein of interest can be tagged with a short 68 amino acid sequence, which is recognized by the E3 ubiquitin ligase TIR1 derived from *Arabidopsis thaliana* in the presence of a small molecule called auxin. Ubiquitination targets the degron-tagged protein for fast degradation by the proteasome. Half times less than 30 minutes have been reported for cytosolic proteins after transferring animals co-expressing a degron-tagged protein and TIR1 on plates containing auxin (Zhang et al., 2015). The AID is therefore faster and more efficient than RNAi. Since AID initiates protein degradation and RNAi initiates mRNA degradation, these two systems do not compete with each other and can be used in parallel.

Here, we identify suppressors of lipid bilayer stress using a novel approach by combining AID and RNAi-based forward genetic screening. Knockdown of MDT-15 by AID was used to induce LBS, which was visualized using the endoplasmic reticulum stress sensors P*atf-4::gfp* and P*hsp-4::gfp*. We screened the RNAi libraries targeting kinases and transcription factors for suppressors. Out of 868 genes, we identified 8 novel hits that robustly blocked LBS caused by MDT-15 knockdown.

## Material and Methods

### Strains

All strains were maintained at 20°C on OP50 *Escherichia coli* as described. Strains used in this study are either available from CGC or upon request:

**IJ1729**: *ieSi57* [P*eft-3::*TIR1::mRuby::*unc-54* 3’UTR; *cb-unc-119*] II; *yh44* [*mdt-15*::degron::EmGFP] III. (Lee et al., 2019)

**SJ4005**: *zcIs4* [P*hsp-4*::GFP] V. (Harding et al., 2000)

**LD1499**: [P*atf-4*(uORF)::GFP::*unc-54*(3’UTR)]

**LSD2096**: *ieSi57* [P*eft-3*::TIR1::mRuby::*unc-54*(3’UTR); *cb-unc-119*] II; *yh44* [*mdt-15*::degron::EmGFP] III; [P*atf-4*(uORF)::GFP::*unc-54*(3’UTR)].

**LSD2102**: *ieSi57* [P*eft-3*::TIR1::mRuby::*unc-54* 3’UTR; *cb-unc-119*] II; *yh44* [*mdt-15*::degron::EmGFP] III; *zcIs4* [P*hsp-4*::GFP] V.

For generation of the screening strain, IJ1729 males was crossed with LD1499. 48 F2 were singled and their offspring were put on plates containing 100 μM Auxin and checked for upregulation of the reporter (Fig. 2a). In parallel, IJ1729 was crossed to SJ4005.

**Figure 2.**
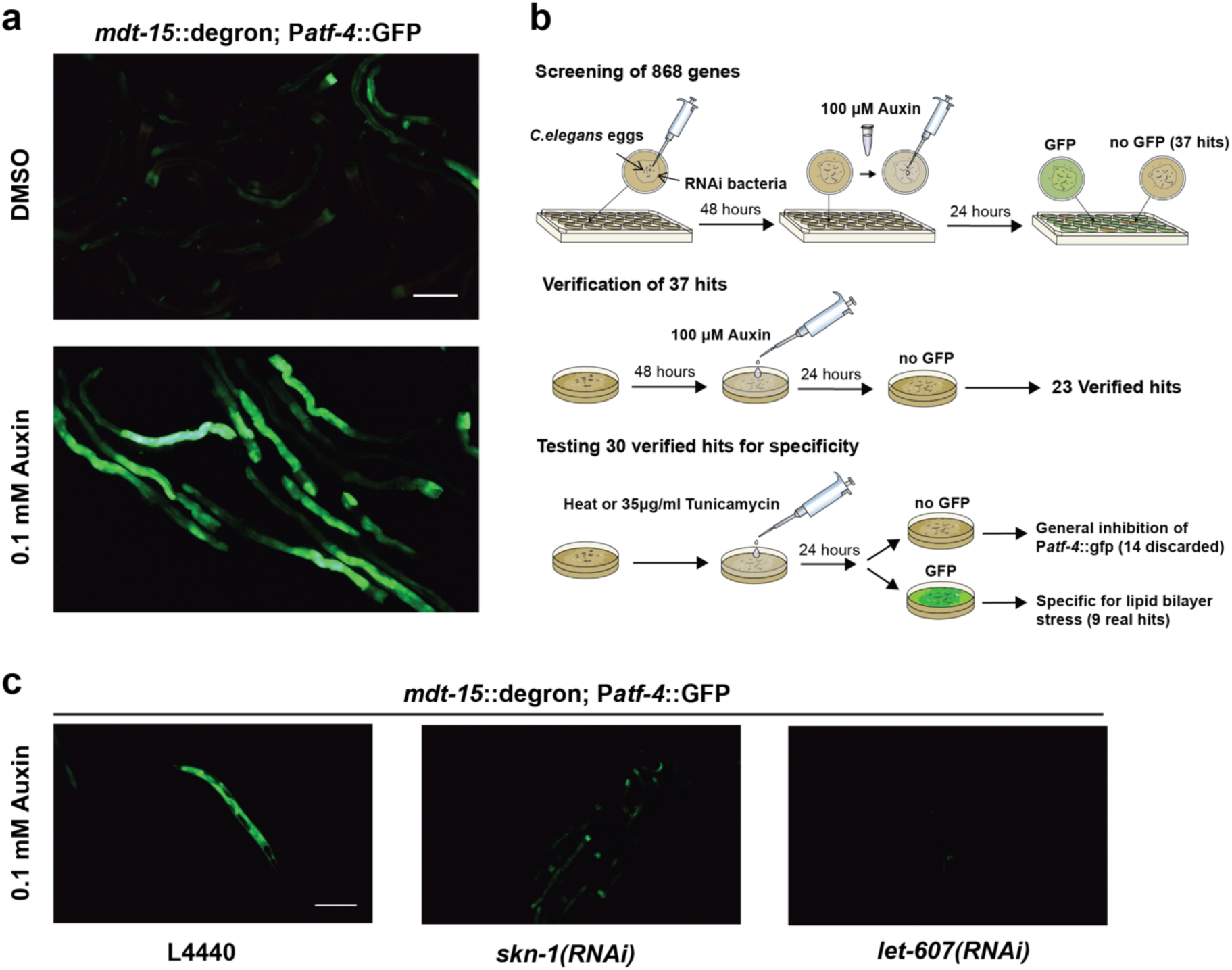
Suppressor screen of LBS. a) Addition of 0.1 mM Auxin degrades *mdt-15*::degron and leads to expression of P*atf-4::gfp* in LSD2096. Pictures were taken 24 h after Auxin addition. Bar = 200 μm b) Summary of the screening outline. c) Addition of 0.1 mM Auxin to degrade *mdt-15*::degron after *skn-1* and *let-60*7 RNAi represses activation of P*atf-4*::gfp in LSD2096. Pictures were taken 24 h after Auxin addition. Bar = 200 μm

### Microscopy

For image acquisition, the animals were put on freshly made 2% agar pads and anesthetized with 1 mM tetramisole. Images were taken with an upright bright field fluorescence microscope (Tritech Research, model: BX-51-F) and a camera of the model DFK 23UX236 (Teuscher and Ewald, 2018).

### Preparation of Auxin

70 mg of Auxin (3-Indoleacetic acid, Sigma #I3750) were dissolved in 10 ml DMSO to yield a 40 mM stock solution and stored at 4°C. The stock was further diluted in M9 to 100 μM before use.

### Suppressor screen design

A detailed step-by-step protocol can be found in the supplement and a schematic outline is shown in Fig. 2b.

Briefly, 24-well plates were filled with NGM containing Ampicillin (100 μg/ml), Tetracyclin (12.5 μg/ml) and 1mM IPTG, seeded with 50 ul of freshly grown RNAi bacteria and dried in a sterile hood. On the next day, plates containing gravid LSD2096 adults were washed off and discarded and the laid eggs were scratched off and collected. Approximately 30-40 eggs were pipetted into a well and incubated at 20°C. After 48 hours, the wells were top coated with 50 μl of 100 μM Auxin, dried in a sterile hood and put at 20°C overnight. The following day, the wells were screened for suppression of the GFP signal. The kinase and transcription factor libraries were screened twice. The preliminary hits had to successfully pass three additional runs on 6 cm plates.

### Heat-shock and tunicamycin treatment

Animals, RNAi bacteria, and plates were prepared as above, without the addition of auxin. Heat-shock was carried out for 1 hour at 37°C, incubated for 5 hours at 25°C, and the animals were checked for GFP expression. Plates were top coated with 0.5 ml of 35 μg/ml tunicamycin (Sigma, T7765), incubated for 6 hours at 25°C, and the animals were checked for GFP expression.

## Results and Discussion

To find a suitable reporter for screening, we used RNAi against *mdt-15* and *fat-6/7* and found two reporters P*atf-4::gfp* and P*hsp-4::gfp* that are activated by LBS (Fig. 1b). For screening, we preferred P*atf-4::gfp* over P*hsp-4::gfp* for its stronger induction of GFP. Crossing *mdt-15(tm2182)* mutant with P*atf-4::gfp* led to a heterogeneous GFP expression, making it impossible to use this strain for screening. We therefore switched to an endogenously degron-tagged *mdt-15* strain (Lee et al., 2019). Unstressed MDT-15::degron *C. elegans* expressed P*atf-4::gfp* only at basal levels at 20°C. Incubation with 100 μM Auxin for 24 hours increased GFP levels drastically and homogenously (Fig. 2a). We additionally observed typical *mdt-15* phenotypes such as small body size, reduced brood size, and a pale appearance (Taubert et al., 2008; Lee et al., 2019). Surprisingly, the eggs of untreated animals were sensitive to bleaching. Either, the screening strain carries a background mutation or degron-tagged *mdt-15* is slightly hypomorph. We continued with our screen by collecting laid eggs off the bacterial lawn.

To gain insights into LBS, we took a targeted RNAi approach. We decided to screen through almost all *C. elegans* kinases (441 genes) and transcription factors (427 genes; Fig. 2b). Our first-pass screening round resulted in 6 kinases and 31 transcription factors (Fig. 2b). To sort out false positives, we tested the preliminary hits on 6 cm plates and ended up with 23 verified hits that block P*atf-4::gfp* induction (Fig. 2b, 2c; Supplementary Table 1). To test whether the hits are specific for lipid bilayer stress, and not general inhibitors of the unfolded protein response, we heat-shocked the animals and treated them with the N-glycosylation-inhibitor tunicamycin (Fig. 2b). Most of the clones not passing this step were wrongly annotated GFP clones (Supplementary Table 1). Reassuringly, we detected *xbp-1*, a transcription factor spliced by IRE-1 (Supplementary Table 1). XPB-1 is known to upregulate *hsp-4* mRNA during UPR (Calfon et al., 2002). To rule out transgene-specific effects, we crossed P*hsp-4::gfp* into *mdt-15::degron*;TIR1 and tested the hits that have passed the previous steps (Fig. 2b). Only the weakest hit, *ztf-1*, did not pass this step (Supplementary Table 1). We ended up with 9 high confidence hits, 8 of them not previously described in *C. elegans* (Table 1). Taken together, with our novel approach of combining AID and RNAi screening, we bypassed developmental and lethal obstacles caused by depletion of *mdt-15*. Our screen revealed a known molecular player (IRE-1), but also identified several new genes important to mount a proper LBS response. Thus, our results provide a proof-of-concept and support the feasibility of combined AID-RNAi screening approaches.

**Table 1.**

| Gene name | Level of repression | Short description |
| --- | --- | --- |
| *drl-1* | Full | Orthologue of mitogen-activated protein kinase kinase kinase 3 isoforms |
| *rpl-14* | Full | Protein of the large ribosomal subunit (60S) |
| *let-607* | Full | Orthologue of the CAMP Responsive Element Binding Protein 3 (CREB3) transcription factor family |
| *gsk-3* | Strong | Orthologue of human GSK3B (glycogen synthase kinase 3 beta) |
| *ire-1* | Strong | Inositol-requiring Enzyme 1, a sensor of the UPR |
| *ntl-4* | Strong | Orthologue of human CCR4-NOT Transcription Complex Subunit 4 (CNOT4) |
| *gtf-2F2* | Strong | Orthologue of human (general transcription factor IIF subunit 2 (GTF2F2) |
| *elt-2* | Strong | Orthologue of the transcription factors GATA4/6 |
| *skn-1* | Strong | Orthologue of the transcription factor Nuclear factor erythroid 2-related factor 2 (NRF2) |

### Regulation of LBS by IRE-1 and XBP-1

We unbiasedly detected IRE-1, which has previously been proposed as a sensor for LBS in yeast and in *C. elegans*, and its target *xbp-1* (Thibault et al., 2012). This confirms the selectivity of our screen. Unfolded proteins in the ER lead to IRE1 oligomerization and the subsequent stimulation of its endoribonuclease activity and splicing of the transcription factor *xbp-1*. However, monomeric IRE1 still displays RNase activity and splices XBP1 mRNA in HeLa cells during LBS (Kitai et al., 2013; Ho et al., 2020). Spliced mRNA of *xbp-1* is much more stable than the unspliced variant. IRE-1 is not needed for heat-shock or tunicamycin-induced UPR, but its target *xbp-1* (Supplemental Table 1). We did not detect *pek-1* and *atf-6*, the other members of the canonical UPR. Thus, we confirmed that IRE-1 branch acts as a major sensor of LBS.

### Immunity response network regulates lipid bilayer stress

Knocking down phosphatidylcholine synthesis leads to activation of genes involved in the immune response (Koh et al., 2018). Many of these transcripts are upregulated in an IRE-1-dependent manner. In addition to *ire-1*, we detected the NRF1,2,3 homologue *skn-1* and the GATA transcription factor *elt-2*. Both are involved in p38-mediated innate immunity (Block et al., 2015; Blackwell et al., 2015). RNAi of *skn-1* was one of our strongest suppressors (Fig. 1c). SKN-1 is a major transcription factor for promoting oxidative stress resistance (Blackwell et al., 2015). There are four isoforms of SKN-1: skn-1a, b, c, d (Blackwell et al., 2015). A previous study demonstrated that IRE-1 has an additional mode of action in its monomeric state: elevated levels of reactive oxygen species leads to sulfenylation of cysteine residues in IRE-1 and activates SKN-1a via the p38 MAPK (Glover-Cutter et al., 2013; Hourihan et al., 2016). Isoform *skn-1a* is similar to mammalian NRF1, which regulates proteostasis and is a transmembrane protein located in the ER (Lehrbach and Ruvkun, 2016; Wang and Chan, 2006). Curiously, *mdt-15* and *skn-1c*, but not *skn-1a*, co-regulate targets involved in detoxification such as *gst-4* (Goh et al., 2014). This suggests that SKN-1a is not only activated by loss of *mdt-15*, but also works independently of MDT-15. *skn-1* knockdown does not only block LBS response, but also reverses the small body size and the small amount of eggs laid (although these eggs do not hatch because *skn-1* knockdown is embryonic lethal). This suggests that some of the observed phenotypes in *mdt-15* mutants or knockdowns are *skn-1*-dependent. The mammalian SKN-1c orthologue NRF2 has been shown to have protective functions during palmitate-induced lipotoxicity in mammalian cells (Cunha et al., 2016; Park et al., 2015). Taken together, this suggests a potential isoform-specific role for SKN-1a during LBS.

*elt-2* is a GATA transcription factor that is essential for the mesodermal cell fate and development of the intestine. While null mutants of *elt-2* are embryonic lethal, post-developmental knockdown shortens lifespan and overexpression extends lifespan (Mann et al., 2016). We observed developmental arrest after *elt-2* knockdown. These arrested larvae were still susceptible to heat and tunicamycin treatments, indicating that the UPR was still intact. Similar to *skn-1, elt-2* is recruited to promoters during *Pseudomonas aeruginosa* infection and co-regulates targets in a p38-mediated fashion (Block et al., 2015). Furthermore, *elt-2* and *mdt-15* cooperate during heavy metal intoxication (Shomer et al., 2019), strengthening the existence of a transcription factor network that cooperatively regulates different stress responses.

### Modulators and activators of the LBS response (let-607, gsk-3 and drl-1)

We found three genes, *let-607, gsk-3*, and *drl-1*, implicated in modulating the ER stress responses. RNAi of *let-607* suppressed the activation of the *atf-4* reporter completely (Fig. 2c). *let-607* is, together with *crh-1* and *crh-2*, one of the CREB3 orthologues in *C. elegans*. The mammalian Creb3 family consists of five members (CREB3/Luman, CREB3L1/OASIS, CREB3L2/BBF2H7, CREB3L3/CREBH, and CREB3L4) and is related to ATF6 and SREBP (Sampieri et al., 2019). All of them are localized in the ER and are similar to ATF6 activated by anterograde transport to the Golgi and subsequent cleavage by S1P or S2P. In humans and mice, CREB3L2 upregulates SEC23 and controls secretary load, especially during bone formation (Saito et al., 2009; Tomoishi et al., 2017; Al-Maskari et al., 2018; Khetchoumian et al., 2019). CREB3 and CREB3L3 are induced after palmitate-induced ER stress and knock down of CREB3 by siRNA sensitizes human islet cells to palmitate-induced ER stress (Cnop et al., 2014). CREB3 has been identified in regulating Golgi-stress and activates ARF4 (Reiling et al., 2013). A previous study in *C. elegans* links *let-607* with the upregulation of *sec-23* and other proteins involved in secretion (Weicksel et al., 2016). *let-607* has also been identified in a screen for suppressors of PolyQ aggregation and suppresses motility defects caused by mutations in the paramyosin ortholog UNC-15, the basement-membrane protein perlecan UNC-52, the myosin-assembly protein UNC-45, and the myosin heavy chain UNC-54 (Silva et al., 2011). In addition, knock down of *let-607* increased expression of cytosolic heat-shock proteins. Based on these previous observations and our results, we propose that *let-607*/CREB3 family is sensing LBS and acts together with the other identified transcription factor encoding genes *xbp-1, skn-1*, and *elt-2* to mount a unique stress response that is different from the canonical UPR.

*drl-1*, also known as *mekk-3*, has been found in a screen looking for enhancers of dauer formation and extends lifespan by simulating dietary restricted-like conditions (Chamoli et al., 2014). Curiously, loss of *drl-1* causes a pale appearance of the animals resembling *fat-6/7* and *mdt-15* mutants, but the mode-of-action seems to be different. The *drl-1* promoter is expressed in vulval muscles, body wall muscles, hypodermis, seam cells, some neurons, and tissues lining the pharynx and anus, but not the intestine. Additionally, knockdown in the intestine using tissue-specific RNAi did not extend lifespan (Chamoli et al., 2014). Knockdown of MDT-15 activates P*atf-4::gfp* and P*hsp-4::gfp* expression mainly in the intestine (Fig 1b). Therefore, knockdown of *drl-1* acts in a cell non-autonomous manner. *drl-1* decreases fat storage by upregulating fatty acid oxidation (Chamoli et al., 2014). The *C. elegans* orthologue of the ribonuclease Regnase-1, *rege-1*, shares many upregulated genes and causes a pale appearance without activation of LBS (Supplementary Table 1; Habacher et al. 2016). This suggests a link between *drl-1* and *rege-1*. However, knockdown of *rege-1* does not phenocopy loss of *drl-1* (Supplementary Table 1). Despite the striking similarities shared by *rege-1* and *drl-1*, only *drl-1* modulates LBS. Intriguingly, *drl-1* knockdown itself causes ER stress at the L2 stage and this mounts a protective effect throughout life (Matai et al., 2019). Since *drl-1* rewires metabolism by mimicking dietary restriction, we speculate that activation of fatty acid oxidation protects from lipotoxicity. Indeed, we observed that starved animals in wells with no food did not upregulate the reporter.

The glycogen synthase kinase-3 (*gsk-3*) has been described as the busiest of all kinases with over 100 targets known, and was found to attenuate palmitate-induced apoptosis (Beurel et al., 2015; Ibrahim et al., 2011). Paradoxically, *gsk-3* inhibits *skn-1* and stabilizes CREB3, both contradicting with the results of our screen (An et al., 2005; Barbosa et al., 2013). If it does not act on the other hits, what could be its mode of action? Activation of the lipid bilayer stress activates autophagy via the IRE-1/XBP-1 axis (Ho et al., 2020). Blocking autophagy in this context causes sickness, sterility and developmental defects. Intriguingly, GSK3 inhibition activates autophagy (Parr et al., 2012). We therefore speculate that prior knock down of GSK3 leads to an elevated rate of autophagy, which protects from LBS and ameliorates the stress response.

### General players in gene expression, but specific for LBS

The last three hits consist of *gtf-2f2, ntl-4*, and *rpl-14*, which are involved in transcription, RNA processing, and translation, respectively. Interestingly, although with RNAi against *gtf-2f2, ntl-4*, and *rpl-14* inactivate general processes, the heat- or tunicamycin induced UPR is still functioning and is not affected. This favors the model that UPR and LBS are differentially regulated (Fig. 3).

**Figure 3.**
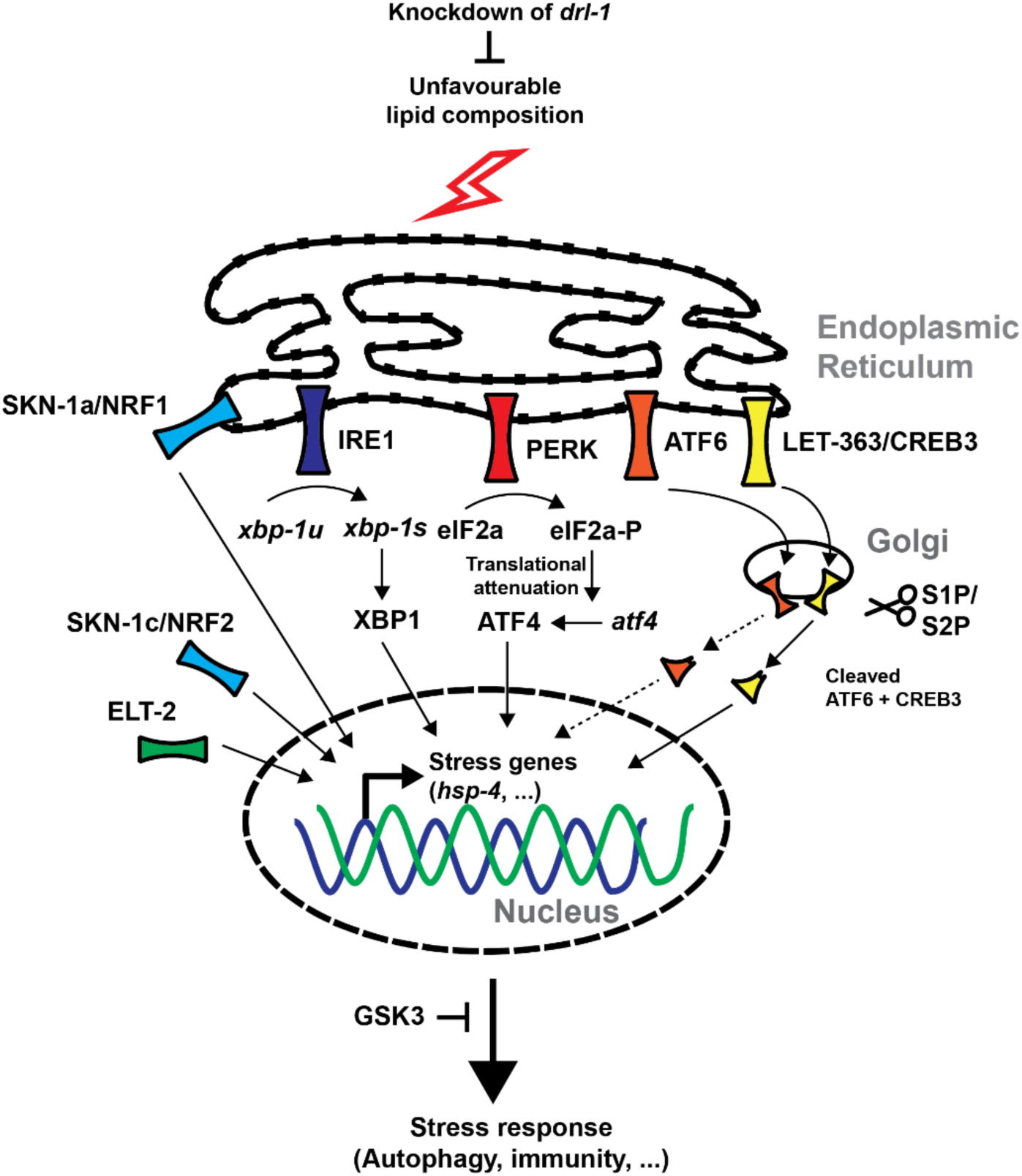
Hypothetical model of LBS in *C. elegans*. Updated model of LBS in *C. elegans* indicates a complex network of transcription factors and up- and downstream modulators. 6 out of 9 our hits are included in this model.

### Summary

We report the identification of 8 novel regulators of the lipid bilayer stress response. We grouped them into three categories (Fig. 3). *skn-1* and *elt-2*, together with the previously characterized *ire-1*, are transcription factors involved in immune response. *let-607* might be activated in parallel with the canonical UPR arms, and *drl-1* and *gsk-3* modulate the ER stress response in our suggested model upstream or downstream, respectively. The last category consists of genes involved in general processes of gene expression. Furthermore, we highlight the potential of combining AID and RNAi-based genetic screens.

## Acknowledgement

We thank Chi Yun and David Ron for the P*atf-4:*:gfp strain, Seung-Jae V. Lee for the *mdt-15::degron* strain, Gary Ruvkun for kinase and transcription factor RNAi libraries, Keith Blackwell and Ewald Lab members for comments on the manuscript. Some strains were provided by the CGC, which is funded by NIH Office of Research Infrastructure Programs (P40 OD010440). Funding from the Swiss National Science Foundation PP00P3_163898 to CYE and ETH Research Foundation Grant ETH-30 16-2 to RV.

## Supplemental files are available at FigShare

Supplementary File1 contains the protocol for 24-well plates AID-RNAi screen. Supplementary Table1 contains the lipotoxicity AID-RNAi screen results.

## References

Acosta-Montaño, P., & García-González, V. (2018). Effects of dietary fatty acids in pancreatic beta cell metabolism, implications in homeostasis. Nutrients, 10(4), 393.

Al-Maskari, M., Care, M. A., Robinson, E., Cocco, M., Tooze, R. M., & Doody, G. M. (2018). Site-1 protease function is essential for the generation of antibody secreting cells and reprogramming for secretory activity. Scientific reports, 8(1), 1–11.

An, J. H., Vranas, K., Lucke, M., Inoue, H., Hisamoto, N., Matsumoto, K., & Blackwell, T. K. (2005). Regulation of the *Caenorhabditis elegans* oxidative stress defense protein SKN-1 by glycogen synthase kinase-3. Proceedings of the National Academy of Sciences, 102(45), 16275–16280.

Barbosa, S., Fasanella, G., Carreira, S., Llarena, M., Fox, R., Barreca, C., … & O’Hare, P. (2013). An Orchestrated Program Regulating Secretory Pathway Genes and Cargos by the Transmembrane Transcription Factor CREB-H. Traffic, 14(4), 382–398.

Beurel, E., Grieco, S. F., & Jope, R. S. (2015). Glycogen synthase kinase-3 (GSK3): regulation, actions, and diseases. Pharmacology & therapeutics, 148, 114–131.

Blackwell, T. K., Steinbaugh, M. J., Hourihan, J. M., Ewald, C. Y., & Isik, M. (2015). SKN-1/Nrf, stress responses, and aging in *Caenorhabditis elegans*. Free Radical Biology and Medicine, 88, 290–301.

Block, D. H., Twumasi-Boateng, K., Kang, H. S., Carlisle, J. A., Hanganu, A., Lai, T. Y. J., & Shapira, M. (2015). The developmental intestinal regulator ELT-2 controls p38-dependent immune responses in adult *C. elegans*. PLoS genetics, 11(5).

Chamoli, M., Singh, A., Malik, Y., & Mukhopadhyay, A. (2014). A novel kinase regulates dietary restriction-mediated longevity in *Caenorhabditis elegans*. Aging Cell, 13(4), 641–655.

Cnop, M., Abdulkarim, B., Bottu, G., Cunha, D. A., Igoillo-Esteve, M., Masini, M., … & Bugliani, M. (2014). RNA sequencing identifies dysregulation of the human pancreatic islet transcriptome by the saturated fatty acid palmitate. Diabetes, 63(6), 1978–1993.

Covino, R., Hummer, G., & Ernst, R. (2018). Integrated functions of membrane property sensors and a hidden side of the unfolded protein response. Molecular cell, 71(3), 458–467.

Cunha, D. A., Cito, M., Carlsson, P. O., Vanderwinden, J. M., Molkentin, J. D., Bugliani, M., … & Cnop, M. (2016). Thrombospondin 1 protects pancreatic β-cells from lipotoxicity via the PERK–NRF2 pathway. Cell Death & Differentiation, 23(12), 1995–2006.

Calfon, M., Zeng, H., Urano, F., Till, J. H., Hubbard, S. R., Harding, H. P., … & Ron, D. (2002). IRE1 couples endoplasmic reticulum load to secretory capacity by processing the XBP-1 mRNA. Nature, 415(6867), 92–96.

Ertunc, M. E., & Hotamisligil, G. S. (2016). Lipid signaling and lipotoxicity in metaflammation: indications for metabolic disease pathogenesis and treatment. Journal of lipid research, 57(12), 2099–2114.

Glover-Cutter, K. M., Lin, S., & Blackwell, T. K. (2013). Integration of the unfolded protein and oxidative stress responses through SKN-1/Nrf. PLoS genetics, 9(9).

Goh, G. Y., Martelli, K. L., Parhar, K. S., Kwong, A. W., Wong, M. A., Mah, A., … & Taubert, S. (2014). The conserved Mediator subunit MDT-15 is required for oxidative stress responses in *Caenorhabditis elegans*. Aging Cell, 13(1), 70–79.

Habacher, C., Guo, Y., Venz, R., Kumari, P., Neagu, A., Gaidatzis, D., … & Ciosk, R. (2016). Ribonuclease-mediated control of body fat. Developmental cell, 39(3), 359–369.

Harding, H. P., Novoa, I., Zhang, Y., Zeng, H., Wek, R., Schapira, M., & Ron, D. (2000). Regulated translation initiation controls stress-induced gene expression in mammalian cells. Molecular cell, 6(5), 1099–1108.

Ho, N., Wu, H., Xu, J., Koh, J. H., Yap, W. S., Goh, W. W. B., … & Thibault, G. (2020). Stress sensor Ire1 deploys a divergent transcriptional program in response to lipid bilayer stress. Journal of Cell Biology, 219 (7): e201909165.

Hou, N. S., Gutschmidt, A., Choi, D. Y., Pather, K., Shi, X., Watts, J. L., … & Taubert, S. (2014). Activation of the endoplasmic reticulum unfolded protein response by lipid disequilibrium without disturbed proteostasis in vivo. Proceedings of the National Academy of Sciences, 111(22), E2271–E2280.

Hourihan, J. M., Mazzeo, L. E. M., Fernández-Cárdenas, L. P., & Blackwell, T. K. (2016). Cysteine sulfenylation directs IRE-1 to activate the SKN-1/Nrf2 antioxidant response. Molecular cell, 63(4), 553–566.

Ibrahim, S. H., Akazawa, Y., Cazanave, S. C., Bronk, S. F., Elmi, N. A., Werneburg, N. W., … & Gores, G. J. (2011). Glycogen synthase kinase-3 (GSK-3) inhibition attenuates hepatocyte lipoapoptosis. Journal of hepatology, 54(4), 765–772.

Koh, J. H., Wang, L., Beaudoin-Chabot, C., & Thibault, G. (2018). Lipid bilayer stress-activated IRE-1 modulates autophagy during endoplasmic reticulum stress. J Cell Sci, 131(22), jcs217992.

Khetchoumian, K., Balsalobre, A., Mayran, A., Christian, H., Chénard, V., St-Pierre, J., & Drouin, J. (2019). Pituitary cell translation and secretory capacities are enhanced cell autonomously by the transcription factor Creb3l2. Nature communications, 10(1), 1–13.

Kitai, Y., Ariyama, H., Kono, N., Oikawa, D., Iwawaki, T., & Arai, H. (2013). Membrane lipid saturation activates IRE 1α without inducing clustering. Genes to Cells, 18(9), 798–809.

Lee, D., An, S. W. A., Jung, Y., Yamaoka, Y., Ryu, Y., Goh, G. Y. S., … & Lee, S.V (2019). MDT-15/MED15 permits longevity at low temperature via enhancing lipidostasis and proteostasis. PLoS biology, 17(8).

Lehrbach, N. J., & Ruvkun, G. (2019). Endoplasmic reticulum-associated SKN-1A/Nrf1 mediates a cytoplasmic unfolded protein response and promotes longevity. Elife, 8, e44425.

Mann, F. G., Van Nostrand, E. L., Friedland, A. E., Liu, X., & Kim, S. K. (2016). Deactivation of the GATA transcription factor ELT-2 is a major driver of normal aging in C. elegans. PLoS genetics, 12(4).

Matai, L., Sarkar, G. C., Chamoli, M., Malik, Y., Kumar, S. S., Rautela, U., … & Mukhopadhyay, A. (2019). Dietary restriction improves proteostasis and increases life span through endoplasmic reticulum hormesis. Proceedings of the National Academy of Sciences, 116(35), 17383–17392.

Min, K., Kang, J., & Lee, J. (2010). A modified feeding RNAi method for simultaneous knock-down of more than one gene in *Caenorhabditis elegans*. Biotechniques, 48(3), 229–232.

Park, J. S., Kang, D. H., & Bae, S. H. (2015). Concerted action of p62 and Nrf2 protects cells from palmitic acid-induced lipotoxicity. Biochemical and biophysical research communications, 466(1), 131–137.

Parr, C., Carzaniga, R., Gentleman, S. M., Van Leuven, F., Walter, J., & Sastre, M. (2012). Glycogen synthase kinase 3 inhibition promotes lysosomal biogenesis and autophagic degradation of the amyloid-β precursor protein. Molecular and cellular biology, 32(21), 4410–4418.

Preston, A. M., Gurisik, E., Bartley, C., Laybutt, D. R., & Biden, T. J. (2009). Reduced endoplasmic reticulum (ER)-to-Golgi protein trafficking contributes to ER stress in lipotoxic mouse beta cells by promoting protein overload. Diabetologia, 52(11), 2369–2373.

Reiling, J. H., Olive, A. J., Sanyal, S., Carette, J. E., Brummelkamp, T. R., Ploegh, H. L., … & Sabatini, D. M. (2013). A CREB3–ARF4 signalling pathway mediates the response to Golgi stress and susceptibility to pathogens. Nature cell biology, 15(12), 1473–1485.

Saito, A., Hino, S. I., Murakami, T., Kanemoto, S., Kondo, S., Saitoh, M., … & Ikawa, M. (2009). Regulation of endoplasmic reticulum stress response by a BBF2H7-mediated Sec23a pathway is essential for chondrogenesis. Nature cell biology, 11(10), 1197–1204.

Sampieri, L., Di Giusto, P., & Alvarez, C. (2019). CREB3 transcription factors: ER-Golgi stress transducers as hubs for cellular homeostasis. Frontiers in Cell and Developmental Biology, 7, 123.

Schwarz, D. S., & Blower, M. D. (2016). The endoplasmic reticulum: structure, function and response to cellular signaling. Cellular and Molecular Life Sciences, 73(1), 79–94.

Shomer, N., Kadhim, A. Z., Grants, J. M., Cheng, X., Alhusari, D., Bhanshali, F., … & Shih, J. (2019). Mediator subunit MDT-15/MED15 and Nuclear Receptor HIZR-1/HNF4 cooperate to regulate toxic metal stress responses in *Caenorhabditis elegans*. PLoS Genetics, 15(12).

Silva, M. C., Fox, S., Beam, M., Thakkar, H., Amaral, M. D., & Morimoto, R. I. (2011). A genetic screening strategy identifies novel regulators of the proteostasis network. PLoS genetics, 7(12).

Singh, J., & Aballay, A. (2017). Endoplasmic reticulum stress caused by lipoprotein accumulation suppresses immunity against bacterial pathogens and contributes to immunosenescence. MBio, 8(3), e00778–17.

Taubert, S., Hansen, M., Van Gilst, M. R., Cooper, S. B., & Yamamoto, K. R. (2008). The Mediator subunit MDT-15 confers metabolic adaptation to ingested material. PLoS genetics, 4(2).

Teuscher, A. C. & Ewald, C. Y. Overcoming Autofluorescence to Assess GFP Expression During Normal Physiology and Aging in Caenorhabditis elegans. BIO-PROTOCOL 8, (2018).

Thibault, G., Shui, G., Kim, W., McAlister, G. C., Ismail, N., Gygi, S. P., … & Ng, D. T. (2012). The membrane stress response buffers lethal effects of lipid disequilibrium by reprogramming the protein homeostasis network. Molecular cell, 48(1), 16–27.

Tomoishi, S., Fukushima, S., Shinohara, K., Katada, T., & Saito, K. (2017). CREB3L2-mediated expression of Sec23A/Sec24D is involved in hepatic stellate cell activation through ER-Golgi transport. Scientific reports, 7(1), 1–11.

Volmer, R., van der Ploeg, K., & Ron, D. (2013). Membrane lipid saturation activates endoplasmic reticulum unfolded protein response transducers through their transmembrane domains. Proceedings of the National Academy of Sciences, 110(12), 4628–4633.

Wang, W., & Chan, J. Y. (2006). Nrf1 is targeted to the endoplasmic reticulum membrane by an N-terminal transmembrane domain Inhibition of nuclear translocation and transacting function. Journal of Biological Chemistry, 281(28), 19676–19687.

Weicksel, S. E., Mahadav, A., Moyle, M., Cipriani, P. G., Kudron, M., Pincus, Z., … & Fernandez, A. G. (2016). A novel small molecule that disrupts a key event during the oocyte-to-embryo transition in *C. elegans*. Development, 143(19), 3540–3548.

Zhang, L., Ward, J. D., Cheng, Z., & Dernburg, A. F. (2015). The auxin-inducible degradation (AID) system enables versatile conditional protein depletion in *C. elegans*. Development, 142(24), 4374–4384.

